# SARS-CoV2 Testing: The Limit of Detection Matters

**DOI:** 10.1101/2020.06.02.131144

**Authors:** Ramy Arnaout, Rose A. Lee, Ghee Rye Lee, Cody Callahan, Christina F. Yen, Kenneth P. Smith, Rohit Arora, James E. Kirby

**Affiliations:** Department of Pathology, Beth Israel Deaconess Medical Center, Boston, MA, USA; Division of Infectious Diseases, Department of Medicine, Beth Israel Deaconess Medical Center, Boston, MA, USA; Harvard Medical School, Boston, MA, USA; Department of Radiology, Beth Israel Deaconess Medical Center, Boston, MA, USA 02215; Department of Surgery, Beth Israel Deaconess Medical Center, Boston, MA, USA 02215; Division of Clinical Informatics, Department of Medicine, Beth Israel Deaconess Medical Center, Boston, MA USA 02215

## Abstract

Resolving the COVID-19 pandemic requires diagnostic testing to determine which individuals are infected and which are not. The current gold standard is to perform RT-PCR on nasopharyngeal samples. Best-in-class assays demonstrate a limit of detection (LoD) of ~100 copies of viral RNA per milliliter of transport media. However, LoDs of currently approved assays vary over 10,000-fold. Assays with higher LoDs will miss more infected patients, resulting in more false negatives. However, the false-negative rate for a given LoD remains unknown. Here we address this question using over 27,500 test results for patients from across our healthcare network tested using the Abbott RealTime SARS-CoV-2 EUA. These results suggest that each 10-fold increase in LoD is expected to increase the false negative rate by 13%, missing an additional one in eight infected patients. The highest LoDs on the market will miss a majority of infected patients, with false negative rates as high as 70%. These results suggest that choice of assay has meaningful clinical and epidemiological consequences. The limit of detection matters.

## Introduction

In response to the SARS-CoV-2 pandemic being declared a public health emergency, clinical and commercial laboratories as well as test kit manufacturers have been submitting diagnostic devices and assays for expedited Emergency Use Authorization by the Food and Drug Administration (FDA EUA). As of June 2020, there were over 85 such EUA issuances for COVID-19 diagnostics (https://www.fda.gov/medical-devices/emergency-situations-medical-devices/emergency-use-authorizations, accessed June 1, 2020). However, optimal use of these assays requires consideration of several issues.

First, NP swabs are generally considered to provide optimal detection early in disease. However, even for this sample type, there is currently no ideal reference standard to establish clinical sensitivities of the available EUA SARS-CoV-2 diagnostic assays (1). Second, details about assay limit of detection (LoD) are often not provided with sufficient detail and transparency to allow facile comparisons. For molecular diagnostic assays, the LoD is generally considered the lowest concentration of target that can be detected in ≥95% of repeat measurements. The LoD is a measure of analytic sensitivity, as opposed to clinical sensitivity, which measures the fraction of infected people detected by a given test. LoDs are sometimes reported in units other than copies of viral genomic RNA per milliliter of transport media (copies/mL), such as TCID_50_, copies/microliter, copies per reaction volume, or molarity of assay target, making comparisons difficult. Third, the LoDs of currently approved EUA nucleic acid amplification and antigen detection tests for SARS-CoV-2 vary up to 10,000 fold (see below) and likely are associated with meaningful differences in clinical sensitivity for these tests. Fourth, although LoDs are quantitative, and RT-PCR tests are inherently quantitative, in practice results for SARS-CoV-2 testing are generally reported qualitatively, as positive or negative, even though viral load may provide both clinically and epidemiologically important information.

Two barriers to quantitative reporting are demonstration that qPCR cycle threshold (Ct) values are repeatable with acceptably low variance and a reliable means of converting from Ct value to viral load. The latter is complicated by a traditional requirement for a standard curve that must span a range of viral loads at least as large as what is observed in the patient population, which can be expensive and time-consuming, especially in a pandemic where the limits of this range are unknown; however, there have been reports demonstrating how appropriate measurements, based on the principles of RT-PCR, can be used as an alternative for reliable conversion of Ct values to viral loads (2, 3).

Here we report on the reliability of Cts for the Abbott SARS-CoV-2 EUA (LoD 100 copies viral RNA/mL transport medium, among the best in class) (4) and a conversion from Ct to viral load, which together support the use of reporting viral loads clinically, and also on an observation based on over 4,700 first-time positive results that makes it possible to estimate the clinical sensitivity and false-negative rate of both this assay and other assays that have received EUA for detecting SARS-CoV-2 infection. These findings have clear implications for patient care, epidemiology, and the social and economic management of the ongoing pandemic.

## Methods

### Setting and time period

All SARS-CoV-2 testing data from The Beth Israel Lahey Health Network from March 26th to May 2nd, 2020 was included in our analysis. The study was deemed exempt by our hospital institutional review board.

### Testing

Tests were performed using the Abbott RealTime SARS-CoV-2 assay, a real-time reverse transcriptase (RT) polymerase chain reaction (PCR) test for qualitative detection of SARS-CoV-2 in NP and oropharyngeal swabs (5). The dual target assay detects both the SARS-CoV-2 RdRp and N genes with a reported LoD of 100 copies/mL. The assay also includes an internal control. Results are reported as positive if the Ct value is ≤31.5, based upon the signal threshold determined by the manufacturer. Ct values for all first-time positive test results were analyzed. Repeat tests were excluded in order to more accurately estimate the range of Ct values of the infected population upon presentation at our medical center. In our internal validation we determined that the LoD with 100% detection for the Abbott m2000 platform was 100 copies/mL (n=80), with Ct mean and standard deviation at this LoD, 26.06±1.03 (4). Note, the Ct determination on Abbott M2000rt platform is alternatively called the fractional cycle number (FCN) and is specifically one way of determining the cycle number at the maximum amplification efficiency inflection point, i.e, the maxRatio, of each amplification curve (6). The FCN has been reported to be a more robust measure for Ct determination than a fixed fluorescence threshold.

### Statistics

Variance was estimated by *R*^2^ of Ct values for repeat tests obtained within 6 hours (n=25 patients, excluding one obvious outlier that by itself accounted for half the total variance: initial Ct 4.4, but repeat negative and attributed to pre-analytic or analytic technical error) and 12 hours (n=51 patients, excluding the same outlier). The conversion from Ct value to viral load was performed using the definition of exponential growth with variable efficiency (2, 3). Efficiency was measured from plots of fluorescence intensity vs. cycle number for 50 positive samples chosen at random, yielding an expression for viral load in copies/mL as a function of Ct (Eq. 6, Supplementary Methods). Per this expression, the expected negative cutoff corresponds to 9.2 copies per mL or ~2 virions per RT-PCR reaction volume (0.5mL), supporting the validity of our parameter estimation.

We used Python (v3.6) and its NumPy, SciPy, Matplotlib, and Pandas libraries to plot linear regression and Theil-Sen slopes with 95% confidence intervals on repeat positives; a normalized cumulative distribution (histogram) of positive results (with reversed x-axis for ease of interpretation); binned histogram by 0.5 log10 units, and linear regression on log10-transformed data.

## Results

Of the 27,098 tests performed on 20,076 patients over the testing period, 6,037 tests were positive (22%), representing 4,774 unique patients. Analysis of repeats within 6 or 12 hours of each other (7) demonstrated high repeatability of Ct values over these short time windows (*R*^2^ 0.70 and 0.63, n=25 and 51, respectively), supporting the validity of this quantitative measure as a basis for assessment of viral load in patients (Fig. 1). We used basic principles of PCR and detailed measurements of PCR efficiency on 50 randomly chosen positive samples to convert from Ct values to viral load, in units of copies of viral RNA per mL of viral transport medium. In order to study the patient population upon presentation without confounding by repeat measurements on the same patients, the remainder of the analysis was on the first positive value for the above 4,774 unique patients.

**Figure 1:**
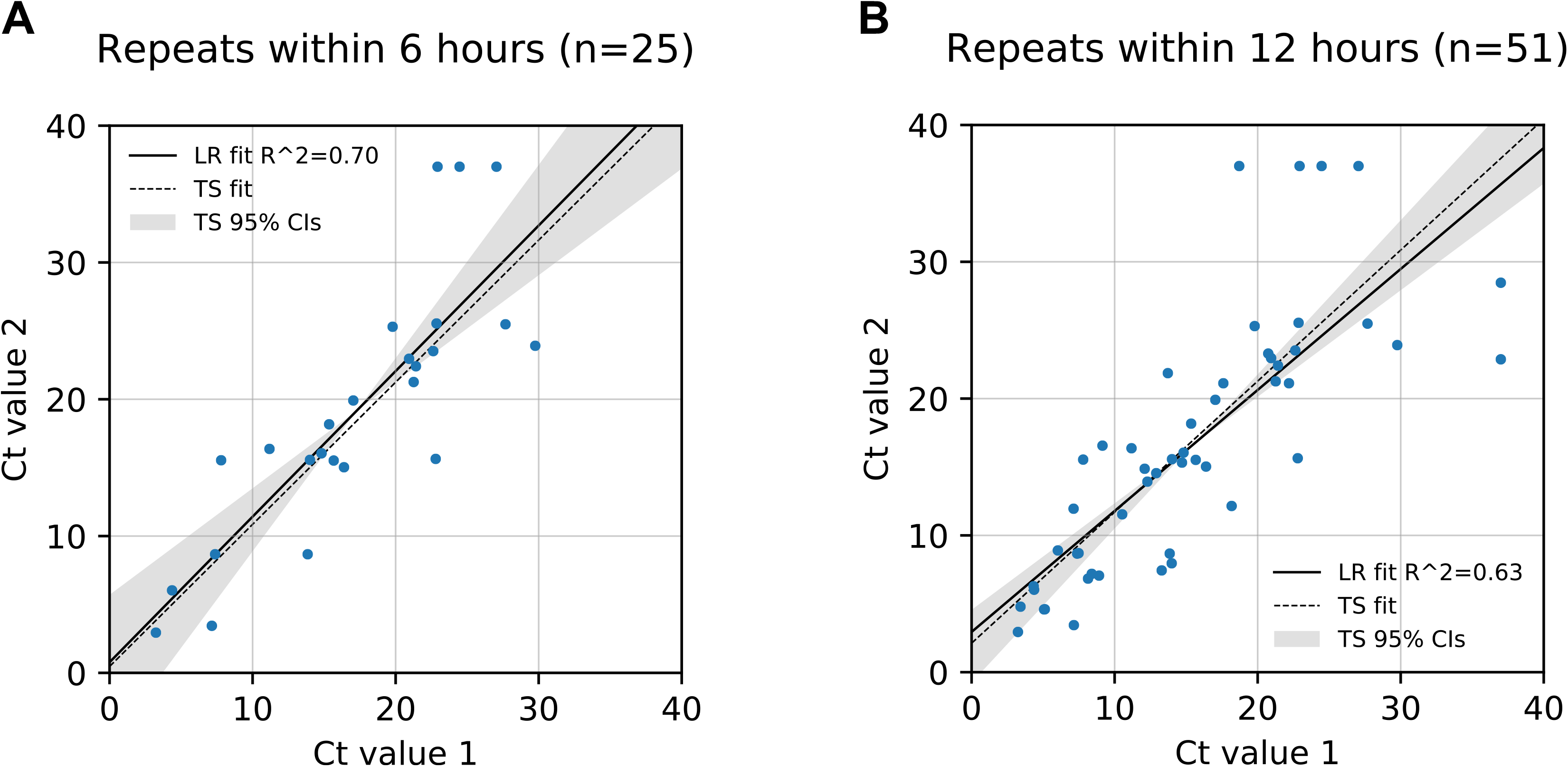
Ct values are highly repeatable. Data points shown are Ct values for SARS-CoV-2 testing of pairs of nasopharyngeal samples obtained within either 6 hours **(A)** or 12 hours **(B)** or each other from the same patient, represented by the X and Y coordinates of each data point. LR = Linear Regression Fit. TS = Theil-Sen Linear Regression Fit. Shade areas indicate 95% confidence interval for TS fit.

Viral loads spanned nearly nine orders of magnitude, from 9 copies/mL to 2.5 billion copies/mL (Fig. 2). Notably, patients were almost equally likely to exhibit low, medium, or high viral loads upon initial testing, with remarkable uniformity down to the LoD of 100 copies/mL (R^2^=0.99). The reason for this uniformity is unknown. Fewer patients had viral loads below the LoD, as reflected by the curve’s departure from the trend in this range. Because the LoD is a 95% confidence limit, the difference between the curve and the trend likely reflects false negatives: the lower the viral load, the greater the likelihood that infection will be missed. By definition, only 5% of patients with viral load at the LoD are expected to be missed (1 in 20 patients); this percentage grows for patients with viral loads below this threshold. Thus, extending the observed trend leftward to the assay’s positive cutoff, which corresponds to approximately two virions per reaction, yields an estimate of the total false negative rate for this assay of 10%, and thus a clinical sensitivity of 90%, or 9 in 10 infected individuals.

**Figure 2:**
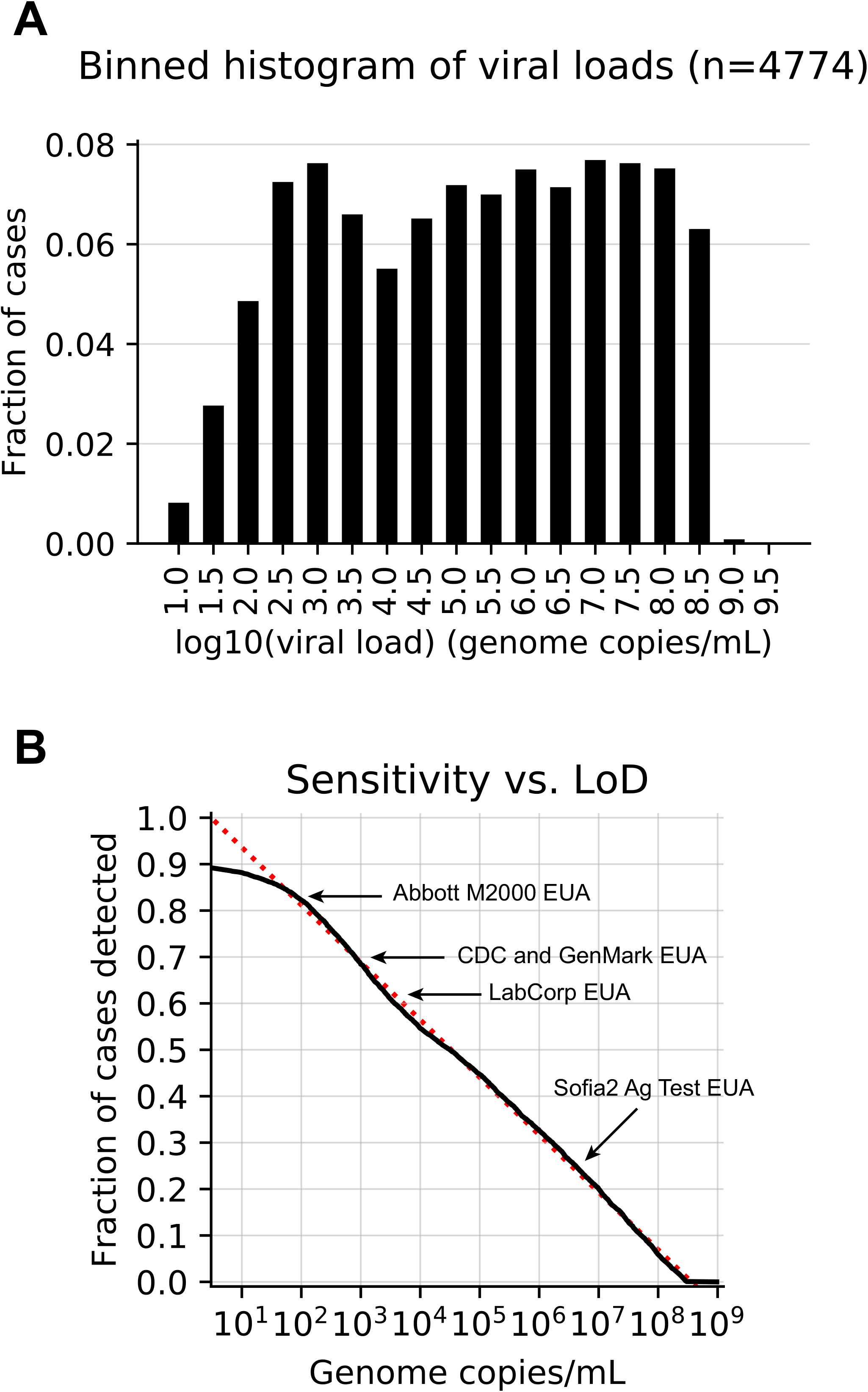
Viral load distribution and LoD. **(A)** Fraction of positive tests binned by 0.5 log10 bins of viral load. **(B)** Cumulative histogram distribution of viral loads showing percent detected as a function of limit of detection - actual, solid line, and trend-line, dotted line.

This method can be used to estimate the clinical sensitivity of assays with other LoDs. For example, an assay with LoD of 1,000 copies/mL, such as that of the CDC assay (8) or Genmark ePlex EUA (9), is expected to detect 77%, or 3 in 4, of infected individuals, for a false-negative rate of 22%. With an LoD of 6,250 copies/mL, the LabCorp COVID-19 RT-PCR EUA test has an estimated clinical sensitivity of 67% and a false-negative rate of 33%, missing approximately 1 in 3 infected individuals. The first EUA antigen detection assay, the Quidel Sofia2 SARS Antigen FIA, has an LoD of approximately 6 million in a contrived universal transport medium sample collection. Although the package insert indicates the LoD using TCID_50_ units, the BEI Resources control material referenced lists both TCID_50_ and genome copies/mL, allowing the calculation of the latter and an associated estimated clinical sensitivity of 31%, i.e., it would miss 7 in 10 infected patients.

## Discussion

The diagnostic priorities in the COVID-19 pandemic are to robustly identify three populations: the infected, the infectious, and the susceptible. Our study addresses the first of these. Specifically, it illustrates the clinical and epidemiologic impact of assay LoD on SAR-CoV-2 diagnosis and the challenges of interpreting and comparing molecular assay results across various platforms. First, viral loads vary widely among infected individuals, from individuals with extremely high viral loads, potential “super-spreaders” who presumably would be picked up by even the least sensitive assays, to those whose viral loads are near, at, or even below the LoD of many assays. Therefore, a substantial fraction of infected patients will be missed by less sensitive assays. Concerningly, some of these missed patients are, have been, or will become infectious, and such misses will undermine public health efforts and put patients and their contacts at risk. This must give pause in the rush to approve additional testing options and increase testing capacity, and emphasizes the importance of defining infectivity as a function of viral load and other factors (e.g. time of exposure), which remains a critical unknown in this pandemic.

Antigen detection assays promise rapid turnaround time, point-of-care implementation, and low cost. For influenza detection, such tests have exhibited substantially lower analytical and clinical sensitivity compared with NAAT tests (10). The poor historical performance for influenza detection led to reclassification of influenza rapid antigen detection tests as Class II devices with a new minimal performance standard of at least 80% sensitivity compared with NAAT (11). Previously, clinical sensitivity of 50-88% for the Quidel Sofia influenza test was noted in several studies in different influenza seasons compared to RT-PCR comparators (12–14). The same trend was observed in our analysis of the single SARS-CoV-2 antigen test introduced thus far with EUA status. Tests with such performance characteristics will identify individuals with the highest viral burden. However, such a high detection threshold will be unlikely to fully meet public or individual health goals in the COVID-19 pandemic.

Our findings also suggest that Ct values and imputed viral loads have clinical utility. Real-time PCR methods in particular are inherently quantitative, and we demonstrate here that they are quite reproducible during repeated clinical sampling over a short time period, with *R*^2^ of 0.70 for repeats within six hours (as a proxy for immediate repeats). We note that because PCR efficiency can fall substantially with PCR cycle number, as we observed here, viral load is ideally calculated not simply as a powers-of-2 transformation of Ct value but based on the observed trend between efficiency and Ct number. This trend may differ by assay: for example, the assay used here includes an internal control whose product may contribute to polymerase inhibition. (This method can be extended to provide confidence limits that incorporate the variance in, e.g., the Ct of the LoD, but this extension is beyond the scope of the current work.)

As yet it is unclear whether or how viral loads affect prognosis, but they at least suggest a measure of infectivity, as well as possibly severity of illness, and, therefore may have value for public health efforts, as we learn which cutoffs may imply minimal or inconsequential infectivity, especially during clearance of infection. We make explicit our assumption that ~2 virions per reaction, translating to a viral load of 9 copies/mL, reflects a 100% detection rate. With stricter cutoffs, clinical sensitivity falls slightly (e.g., from 90% to 86% for an assay with an LoD of 100 copies/mL, if using a cutoff of 4 copies/mL, or a single virion per reaction, and to 79% if using a cutoff of 0.7 copies/mL, or a single virion per 3mL transport tube). Regardless, these different assumptions have essentially no effect on the relative clinical sensitivities of different assays. While it is theoretically possible that even lower levels of infection are possible, making our estimates of clinical sensitivity upper limits, we believe potential for contagion at these levels is highly unlikely, as that would assume that breathing, a cough, or a sneeze would transmit more particles than can be obtained by dedicated and vigorous physical swabbing of the actual nasopharynx.

To control the pandemic, ultimately we will need diagnostics for all three populations of interest, infected, infectious, and susceptible, and for that we will need to understand whether and how viral load relates to infectiousness. As we have shown, assays with higher LoD are likely to miss non-negligible fractions of infected individuals. However, individuals with viral burdens low enough to be missed by some assays may prove to be less infectious. *In vitro*, approximately only 1 of 10,000 genome copies in viral cultures may be associated with a tissue culture infectious viral particle based on standard preparation such as BEI Resources NR-52866(15). However, it is unclear how or whether this fraction might change with viral load for patients *in vivo*.

The ultimate lesson from these studies bears repetition: LoD matters and directly impacts efforts to identify, control, and contain outbreaks during this pandemic. Various assays report out LoDs in manners that are often difficult to comprehend, for example, TCID_50_ values that may related to viral copy numbers in different ways depending on the viral preparation, or units of copies/μL (1 copy/μL = 1,000 copies/mL) or attomolar quantities (1 attomolar = 602 copies/mL). We therefore suggest that viral copies/mL be used as a universal standard metric, so that cross comparison between assays can readily be made. It is clear that viral load matters, and therefore LoD values should be readily evaluable and in the public domain.

## Acknowledgements

K.P.S. was supported by the National Institute of Allergy and Infectious Diseases of the National Institutes of Health under award number F32 AI124590. The content is solely the responsibility of the authors and does not necessarily represent the official views of the National Institutes of Health. We would like to thank the clinical laboratory scientists and volunteers in the Beth Israel Deaconess Medical Center microbiology laboratory for generating the data used in this manuscript.

